# Increasing role of pyrethroid-resistant *Anopheles funestus* in malaria transmission in the Lake Zone, Tanzania: implications for the evaluation of novel vector control products

**DOI:** 10.1101/2021.04.01.438043

**Authors:** Nancy S. Matowo, Jackline Martin, Manisha A. Kulkarni, Jacklin F. Mosha, Eliud Lukole, Gladness Isaya, Boniface Shirima, Robert Kaaya, Catherine Moyes, Penelope A. Hancock, Mark Rowland, Alphaxard Manjurano, Franklin W Mosha, Natacha Protopopoff, Louisa A. Messenger

## Abstract

*Anopheles funestus* is playing an increasing role in malaria transmission in parts of sub-Saharan Africa, where *An. gambiae* s.s. has been effectively controlled by long-lasting insecticidal nets. We investigated vector population bionomics, insecticide resistance and malaria transmission dynamics in 86 study clusters in North-West Tanzania. *An. funestus* s.l. represented 94.5% (4740/5016) of all vectors and was responsible for the majority of malaria transmission (96.5%), with a sporozoite rate of 3.4% and average monthly entomological inoculation rate (EIR) of 4.57 per house. Micro-geographical heterogeneity in species composition, abundance and transmission was observed across the study district in relation to key ecological differences between northern and southern clusters, with significantly higher densities, proportions and EIR of *An. funestus* s.l. collected from the south. *An. gambiae* s.l. (5.5%) density, principally *An. arabiensis* (81.1%) and *An. gambiae* s.s. (18.9%), was much lower and closely correlated with seasonal rainfall. Both *An. funestus* s.l. and *An. gambiae* s.l. were similarly resistant to alpha-cypermethrin and permethrin. Overexpression of *CYP9K1, CYP6P3, CYP6P4* and *CYP6M2* and high L1014F-*kdr* mutation frequency were detected in *An. gambiae* s.s. populations. Study findings highlight the urgent need for novel vector control tools to tackle persistent malaria transmission in the Lake Region of Tanzania.

## Introduction

The widespread deployment of primary vector control interventions, principally long-lasting insecticidal nets (LLINs) and indoor residual spraying (IRS), has substantially reduced malaria incidence across sub-Saharan Africa [1,2]. Between 2000 and 2015, 68% of the 1.5 billion malaria cases averted can be attributed to LLINs alone [1]. However, current estimates indicate the rates of decline have begun to stagnate [2]. Tanzania is among the 10 sub-Saharan African countries where malaria burden is concentrated [3], contributing to 5% of global malaria deaths [2]. Malaria infection varies nationwide with an average prevalence of 7.3% in children under 5 years of age in 2017 [4]. Vector control by the National Malaria Control Programme (NMCP) is based on sustaining high LLIN access and use [5], via universal coverage campaigns supplemented with continuous distribution from school net programmes, antenatal care campaigns and the expanded programme for immunization; and targeted IRS in high transmission areas in the North-West [6]. Effective and sustainable malaria vector control is plagued by a number of challenges, including the evolution of vector behavioural and physiological resistance to current control interventions [7]. In the majority of sentinel districts across Tanzania, *Anopheles* mosquitoes have demonstrated reduced susceptibly to at least one public health insecticide [8,9].

Continued use of insecticide-based malaria control tools has been linked with changes in *Anopheles* feeding and resting behaviors and relative species composition [10-13]. In some countries, *Anopheles funestus* sensu stricto (s.s.) has historically played a significant role in malaria transmission [14-17] largely due to its predominantly anthropophilic and endophilic tendencies [18], intense pyrethroid resistance [19-24] and greater daily survival probabilities (higher parity rates) [25,26]. In other areas, notably south-east Tanzania [25,27], far north-west Tanzania [28] and parts of Kenya [13], this species is rapidly replacing *An. gambiae* s.s. and *An. arabiensis*, following the scale-up of vector control interventions, and has been found with some of the highest *Plasmodium* sporozoite rates [25]. Increasing *An. funestus* population densities and vectorial capacity in these regions may be due to recent escalations in pyrethroid resistance intensities [13,25,27], but also changes in aquatic larval habitats which are more permissible for *An. funestus* breeding [29].

Malaria prevalence around Lake Victoria remains amongst the highest in Tanzania [30], despite high community-level coverage with LLINs, and periodic IRS campaigns [6,28]. Factors driving persistent malaria transmission in the region, including the relative importance of *An. funestus* sensu lato (s.l.) as a major vector species, are poorly characterised but warrant investigation for the design and strategic deployment of new vector control tools. We assessed vector population bionomics, malaria transmission dynamics, phenotypic insecticide resistance and underlying molecular and metabolic resistance mechanisms in 86 study clusters in Misungwi district, north-west Tanzania, prior to a randomised controlled trial assessing the efficacy of next-generation LLINs to improve malaria control [31].

## Results

### Household characteristics

A total of 1,593 households were visited during two cross-sectional entomological field surveys, across 86 clusters in Misungwi district, North-West Tanzania on the southern shore of Lake Victoria, between August and December 2018 (Figure 1A).

**Figure 1:**
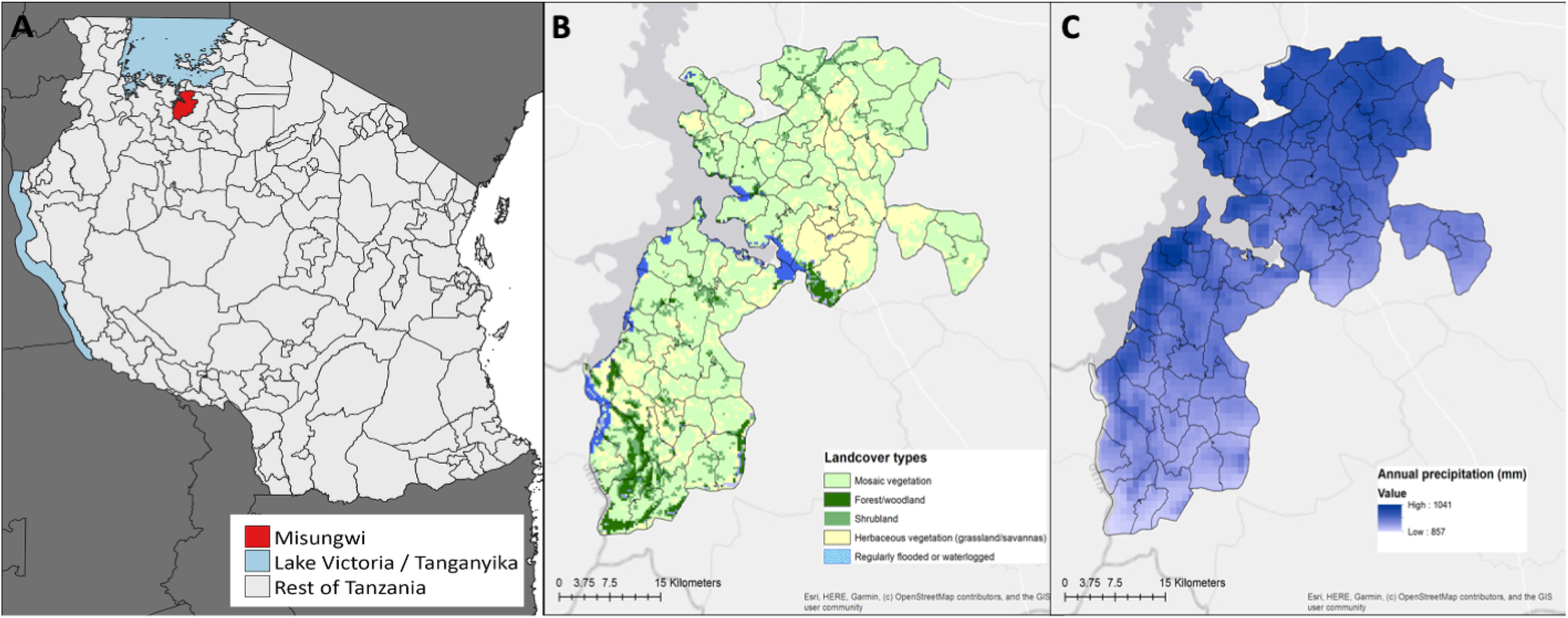
Study area in Misungwi district, north western Tanzania, displaying A: location of Misungwi in the Lake Region; B: landcover features of study clusters; and C: annual precipitation (mm) in study clusters.

**Figure 2:**
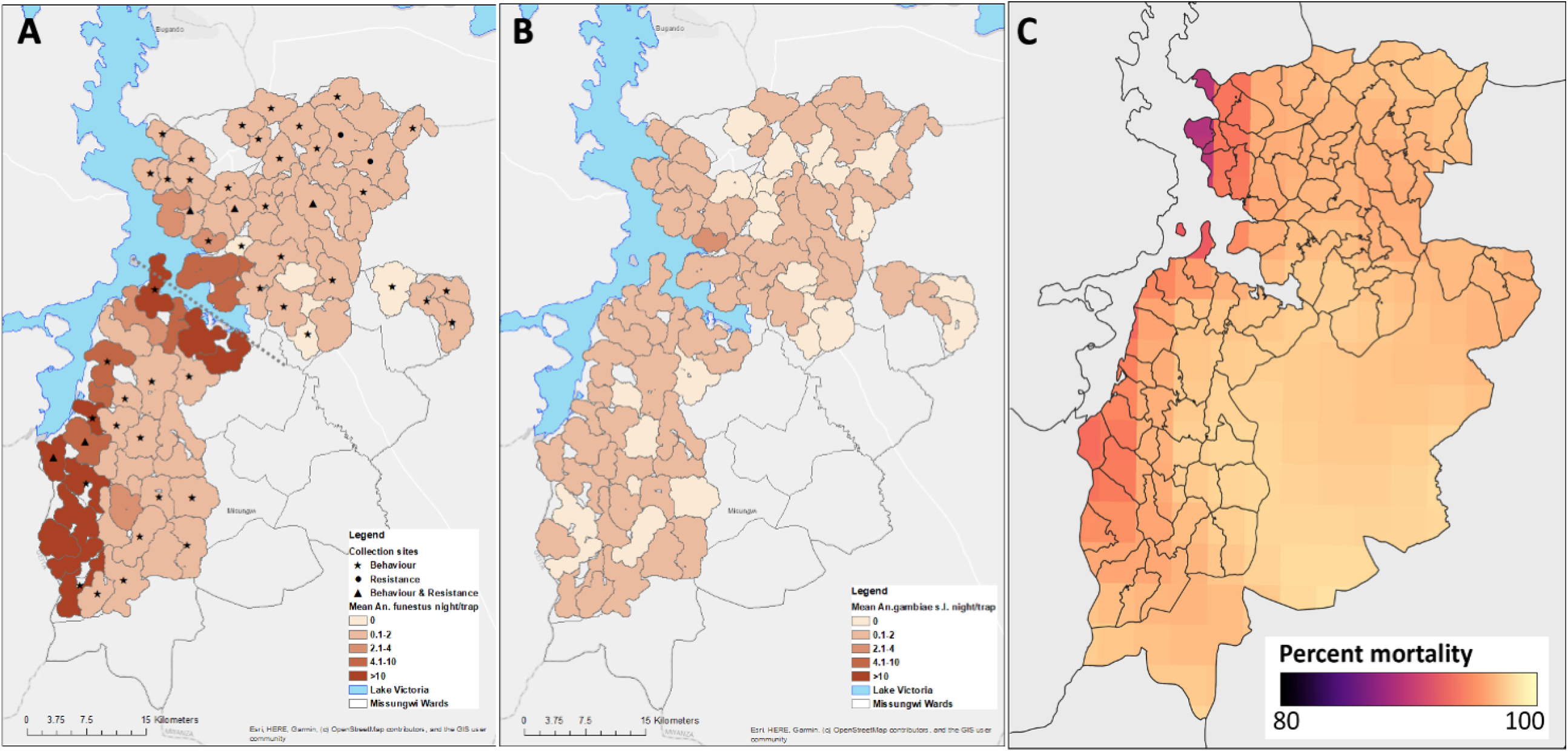
Study area in Misungwi district, north western Tanzania, displaying A: distribution of *Anopheles funestus* s.l. and collection methods per cluster (hashed line indicates delineation between northern and southern clusters); B: distribution of *Anopheles gambiae* s.l.; and C: predicted pyrethroid resistance *for An. gambiae* s.l. (mean percentage mortality).

**Table 1.** Household characteristics in the study area in Northern and Southern clusters. The district spans two agro-ecological zones, based on vegetation land cover and rainfall (Figure 1), that are divided roughly into northern and southern clusters.

| Outcome | All |  | Northern villages |  | Southern villages |  |
| --- | --- | --- | --- | --- | --- | --- |
|  | N | % /mean [95% CI] | N | %/ mean [95% CI] | N | %/mean [95% CI] |
| Total HHs visited | 1593 | N/A | 980 | N/A | 613 | N/A |
| Total consents given | 1373 | N/A | 847 | N/A | 526 | N/A |
| Total HHs analysed | 1372 | N/A | 846 | N/A | 526 | N/A |
| Average number of people per household | 1372 | 6.6 [6.4 - 6.8] | 846 | 6.5 [6.2-6.8] | 526 | 6.8 [6.4-7.1] |
| Average number of sleeping space | 1372 | 2.7 [2.6 - 2.8] | 846 | 2.7 [2.6 -2.8] | 526 | 2.7 [2.5 -2.8] |
| Average altitude (meters) | 1372 | 1194.9 [1187.5 -1202.4] | 846 | 1195.3 [1186.2-1204.3] | 525 | 1194.4 [1181.0-1207.8] |
| % household with iron roof | 1372 | 67.4% [63.2 -71.2] | 846 | 67.2% [61.0 - 72.7] | 526 | 67.7% [62.5 -72.5] |
| % household with open eaves | 1372 | 40.6% [37.0 - 44.3] | 846 | 38.4% [33.7 -43.4] | 526 | 44.1% [38.6 -49.8] |
| % household with screened windows | 1372 | 29.9% [26.2-33.7] | 846 | 33.1% [28.1 -38.5] | 526 | 24.5% [20.1-29.5] |
| % houses made of brick walls | 1372 | 67.5% [62.5 - 72.1] | 846 | 68.9% [62.6 - 74.6] | 526 | 65.2% [56.6- 72.9] |
| % houses with no ceiling | 1372 | 97.2% [95.4 - 98.3] | 846 | 96.3% [93.5 - 98.0] | 526 | 98.5% [96.9 - 99.3] |
| % houses with modern constructed materials | 1372 | 51.8% [ 47.8-55.8] | 846 | 53.7% [48.1-59.1] | 526 | 48.9% [43.1-54.6] |
| % household owning cattle and goats | 1372 | 43.8% [39.8 - 47.9] | 846 | 41.3% [36.0 -46.8] | 526 | 47.7% [41.8 -53.7] |
| % of household owning at least one ITN | 1372 | 94.9% [93.5 -96.0] | 846 | 95.3% [93.5 -96.6] | 526 | 94.3% [91.5 -96.2] |
| Mean number of ITN per house | 1372 | 2.3 [2.2-2.4] | 846 | 2.4 [2.2-2.5] | 526 | 2.1 [2.0 -2.3] |
| Population access to ITN (One net for every two people) | 1372 | 67.9% [65.8 - 70.1] | 846 | 70.4% [67.8-73.1] | 526 | 63.9% [60.5 - 67.3] |
| % HHs with enough nets to cover their sleeping places | 1372 | 62.3% [59.0-65.7] | 846 | 64.1% [59.9-68.2] | 526 | 59.5% [53.7-65.4] |
| % of household sprayed in 2015 | 1372 | 54.2% [49.8-58.5] | 445 | 52.5% [46.5 -58.4] | 299 | 56.8% [50.3- 63.2] |

HH; household; ITN; insecticide treated net; N/A; not applicable

The study had an overall response rate of 86.2% (1372/1592), with consent to participate in the survey given from an adult/head of the household. Ten per cent (164) of dwellings were found vacant, 1.0% (16) were not located, 0.2% (3) not visited due to accessibility and 0.1% (2) were ineligible (no children under 15 years) during the survey period. A small proportion 2.2% (35) refused to participate in the study. The average altitude of study households was 1194.9 meters above sea level (Table 1). Similar proportions of houses were classified as improved or unimproved, based on construction with modern or traditional materials, respectively (Table 1; Figure 3A). Notably, few houses had mosquito proofing materials over the windows (29.9%) and almost no houses had ceilings (97.2%); 40.6% of houses had open eaves (Table 1; Figure 3A and B). The average household size was 6.6 persons and mean number of room/sleeping place was 2.7 per house. Forty-four per cent of households owned at least one livestock (mostly goats and cattle), which were usually kept outdoors about 20 metres away from the house.

**Figure 3:**
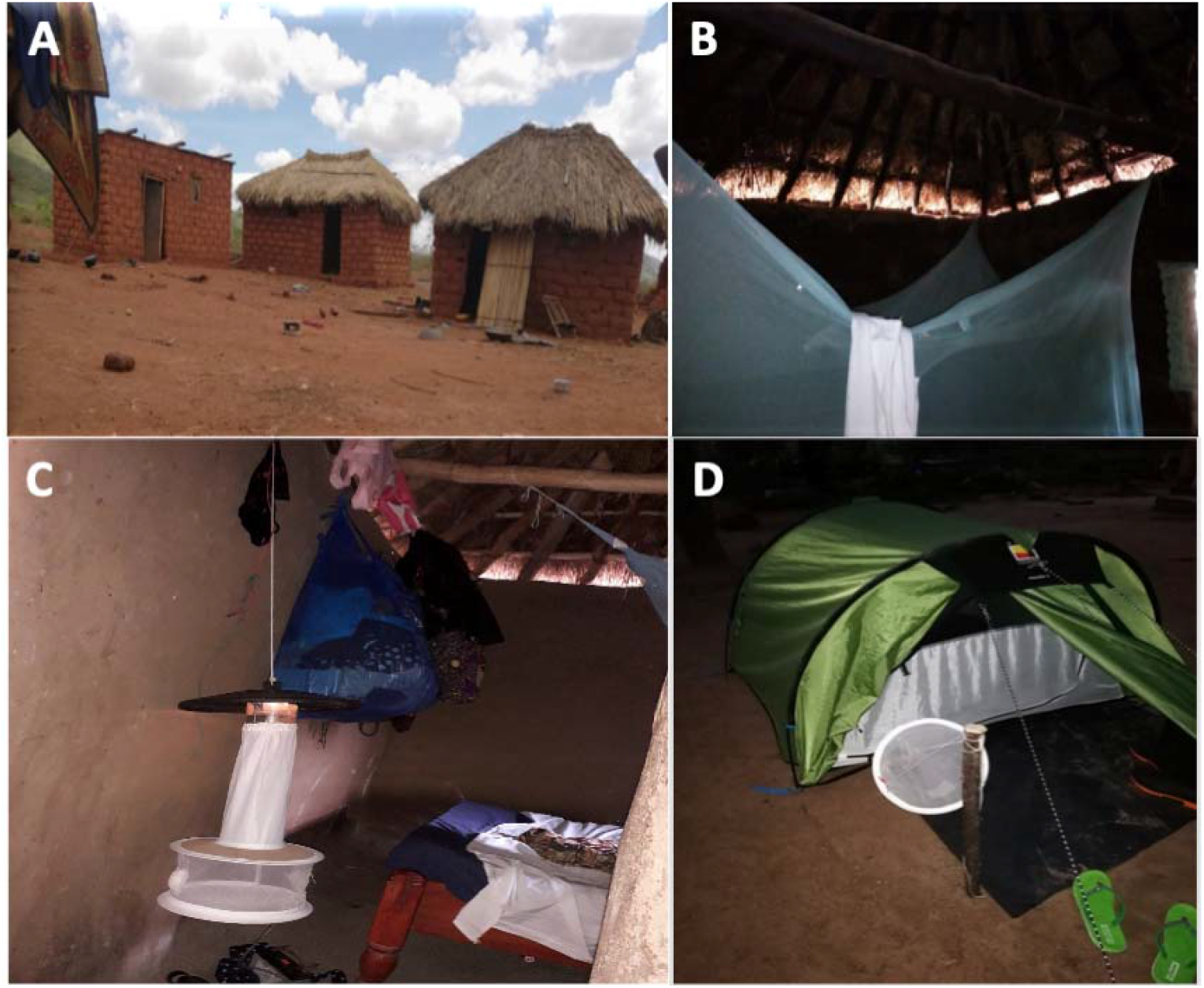
A examples of different traditional house constructions, using local materials. B: the inside of a typical house with open eaves. C: a CDC-LT hung at the base of a sleeping space for sampling mosquitoes indoors. D: a Furvela tent trap set up for catching host-seeking female *Anopheles* mosquitoes outdoors.

LLIN ownership was very high in the study area with the majority of families owning at least one LLIN (94.9%); LLIN access was comparatively lower, however, the majority of households had enough LLINs to cover all of their sleeping places (62.3%). About 54.2% of households were sprayed during the 2015 IRS National Malaria Control Campaign (Table 1). There were no significant differences in household characteristics, including size, altitude, construction materials, however population access to insecticide-treated net (ITN) access was slightly higher in northern than southern clusters (Table 1).

### Vector distribution, species composition, relative abundance and seasonality

A total of 23,081 mosquitoes, comprising 23.1% (5329) Anophelines and 76.9% (17,752) Culicines, were collected using Centers for Disease Control and Prevention light traps (CDC-LTs) during two cross-sectional survey rounds between August and December 2018, for a total of 1373 trap nights (Figure 3C). Most mosquito collections (82.1%) did not experience rainfall and 35.6% of collections had moderate winds.

Of the Anophelines collected, 94.1% (5016) were malaria vectors comprised of 94.5% (4740) *An. funestus* s.l. and 5.5% (276) *An. gambiae* s.l. Significantly greater numbers of *An. funestus* s.l. were collected across the study area compared to *An. gambiae* s.l., p<0.001 (average number of mosquitoes caught per trap per house per night were: *An. gambiae* s.l = 0.20 [95% CI: 0.15-0.27], *An. funestus* s.l = 3.45 [95% CI: 2.58-4.32]) (Table 2). Within the *An. funestus* complex, the predominant species found was *An. funestus* s.s. (92.9%; 710/764 selected for species-specific PCR); other species identified were *An. parensis* (6.5%) and *An. rivulorum* (0.5%). Of the 194 *An. gambiae* s.l selected for sibling species identification, 81.1% were *An. arabiensis* and 18.9% were *An. gambiae* s.s (Table 2).

**Table 2.** Malaria vector species composition, sporozoite rate and entomological inoculation rate (EIR) per study zone.

| Outcome | Overall |  | Northern villages |  | Southern villages |  |
| --- | --- | --- | --- | --- | --- | --- |
|  | N | %/ mean [95% CI] | N | % /mean [95% CI] | N | % /mean [95% CI] |
| Total HH/night collection | 1372 | N/A | 846 | N/A | 526 | N/A |
| Mean mosquitoes per night per house | 23,081 | 16.8 [11.6-22.0] | 9830 | 11.6 [7.5-15.8] | 13251 | 25.2 [13.9-36.4] |
| Mean *Anopheles* vectors | 5016 | 3.7 [1.9 - 5.4] | 788 | 0.9 [0.5-1.4] | 4228 | 8.0 [3.9 -12.1] |
| Total *An. gambiae* s.l | 276 | 0.2 [0.1-0.3] | 176 | 0.2 [0.1-0.3] | 100 | 0.2 [0.1-0.3] |
| Total *An. funestus* s.l | 4740 | 3.5 [1.7-5.2] | 612 | 0.7 [0.3-1.1] | 4128 | 7.8 [3.8 - 11.9] |
| Total *Culex* species | 15609 | 11.4 [7.7-15.0] | 7417 | 8.8 [5.6-12.0] | 8192 | 15.6 [7.8-23.3] |
| Proportion of *An. arabiensis*, n/N | 150/185 | 81.1% [67.9 -89.7] | 70/103 | 68.0% [49.6 - 82.1] | 80/82 | 97.6% [91.8- 99.3] |
| Proportion of *An. gambiae* s.s, n/N | 35/185 | 18.9% [10.3-32.1] | 33/103 | 32.0% [17.9 - 50.4] | 2/82 | 2.4% [0.7- 8.2] |
| Proportion of *An. funestus* s.s, n/N | 710/764 | 92.9% [89.7- 95.2] | 202/220 | 91.8% [85.9 - 95.4] | 508/544 | 93.3% [89.1- 96.0] |
| Sporozoite rate, n/N | 67/1963 | 3.4% [2.5- 4.6] | 13/603 | 2.2% [1.1 - 4.1] | 54/1360 | 4.0% [2.9 - 5.4] |
| Monthly EIR/house | 1349 | 4.4 [1.2- 7.7] | 831 | 0.6 [0.1 - 1.2] | 518 | 9.6 [2.7- 16.4] |

EIR: entomological inoculation rate; HH: household; N/A: not applicable

Overall, significantly higher mosquito densities were observed in villages located in the southern clusters compared to the northern clusters (average number of *Anopheles* caught per trap per house per night in the northern zone=0.93, southern zone=8.04; Density Ratio (DR)=6.09 [95% CI: 3.00-12.38]; p<0.001) (Table 2). There were significantly more *An. funestus* s.l. sampled from households in the southern part of the study area (average/house/night=7.85), compared to the north (0.72; DR=7.92 [95% CI: 3.76-16.67]; p<0.001). However, there was no statistical difference in *An. gambiae* s.l. collected between the two locations with an average of 0.21 per night in the northern zone and 0.19 in the southern zone (DR=1.28 [95% CI: 0.69-2.38]; p=0.431) (Table 2).

Amongst sibling species of the *An. gambiae* complex, there were marked spatial and seasonal fluctuations. Most *An. gambiae* s.s (94.3%) were collected from the northern zone and more than 71.5% of *An. funestus* s.s from the southern part. Both *An. funestus* s.s and *An. arabiensis* predominated throughout the study period, but *An. gambiae* s.s. abundance peaked in December in the middle of the rainy season. An analysis of bioclimatic and landcover characteristics across the study area demonstrated several ecological differences between the northern and southern zones, with the former composed mostly of grassland and cropland (91%), with smaller proportions of shrubland and forest (7%) and areas prone to regularly flooding (1%); and the latter with less grassland and cropland (80%) and greater proportions of shrubland and forest (14%) and areas prone to regular flooding (3%) (Supplementary table 1). Furthermore, differences in rainfall were also observed between the northern and southern zones, with villages in the north receiving slightly higher average annual precipitation (959.5mm) than in the south (911.5mm).

While overall *An. gambiae* s.l density was low, it was closely correlated with seasonal rainfall patterns. Mean *An. gambiae* s.l caught per house during the dry season (August and September; average precipitation of 5-7 mm) was 0.14 but rose significantly by two-fold (DR=1.73 [95% CI: 1.08-2.78]; p=0.02) in the wet season (October, November and December; average precipitation of 147.3-158.5 mm). By comparison, the highest *An. funestus* s.l densities were observed during the dry months (mean=4.61) (Table 3).

The majority of sleeping spaces/beds where the CDC-LTs were installed had either Olyset® (60.5%; 830/1372) or PermaNet® 2.0 LLINs (36.0%; 494/1372); 0.5% (7) had both Olyset® and PermaNet® LLINs, previously distributed through mass universal replacement campaigns (URCs) that was conducted between 2014 and 2017 to achieve universal coverage [32]. The remaining 2.6% (48) of nets had no labels and 0.4% (6) were missing data on net type. There was no significant difference in malaria vector densities between rooms with the two main types of LLIN (average number of malaria vectors per house per night with Olyset® LLINs=3.59 [95% CI: 1.64 -5.54], *versus* PermaNet® 2.0 LLINs=3.89 [95% CI: 1.78 -6.00]; p=0.111).

### *Plasmodium falciparum* infection and entomological inoculation rate

A total of 1963 *Anopheles* mosquitoes (603 and 1360 from the northern and southern clusters, respectively) were tested for the presence of *Plasmodium falciparum* circumsporozoite protein (CSP), with 67 found infected, giving an overall sporozoite rate of 3.4% [95% CI: 2.5-4.6] (Table 2). Of the *An. gambiae* s.l. and *An. funestus* s.l. individuals which tested CSP positive, 6.1% (4/66) were *An. gambiae* s.s., 1.5% (1/66) *An. arabiensis*, and 77.3% (51/66) *An. funestus* s.s., respectively; the remaining samples could not be amplified by PCR. Sporozoite rates were similar in *An. funestus* s.l. (3.44%) compared to *An. gambiae* s.l (3.21%) (Table 3).

**Table 3.**
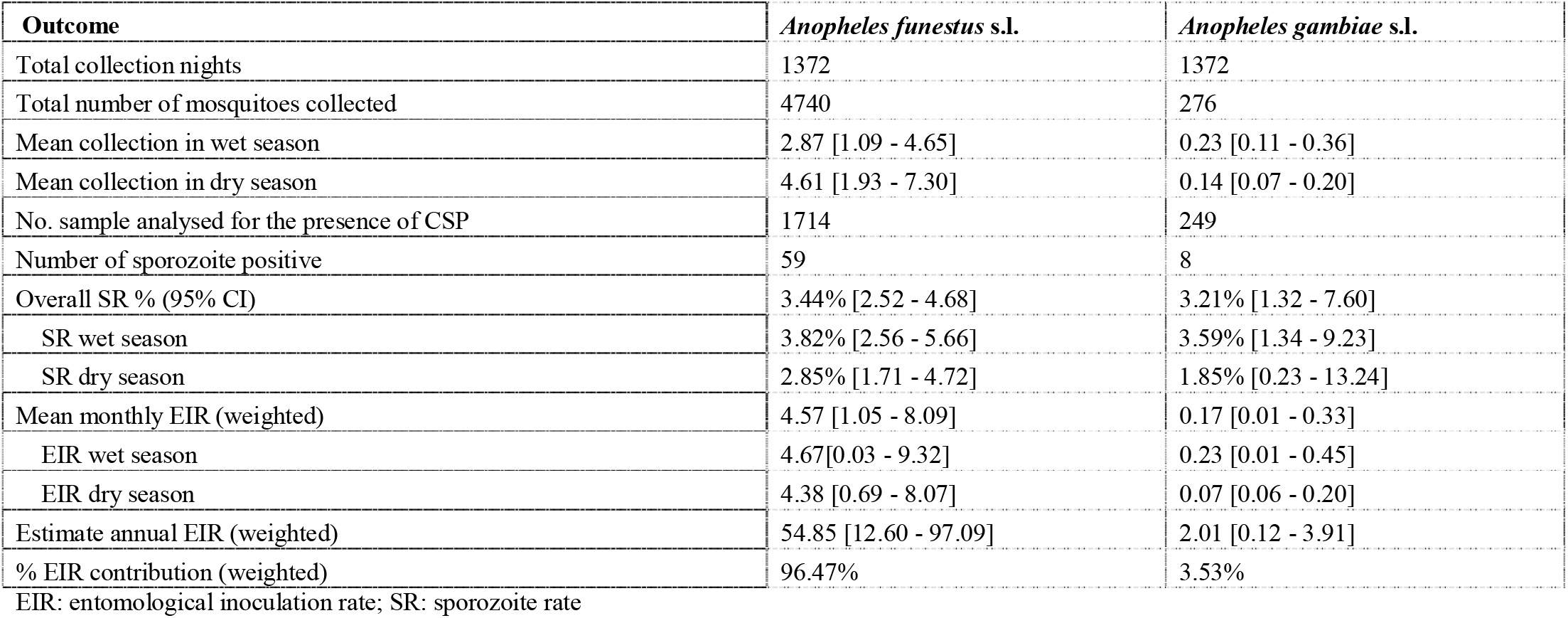
Seasonal variation between *An. funestus* s.l. and *An. gambiae* s.l. sporozoite rate and entomological inoculation rate (EIR).

| Outcome | *Anopheles funestus* s.l. | *Anopheles gambiae* s.l. |
| --- | --- | --- |
| Total collection nights | 1372 | 1372 |
| Total number of mosquitoes collected | 4740 | 276 |
| Mean collection in wet season | 2.87 [1.09 - 4.65] | 0.23 [0.11 - 0.36] |
| Mean collection in dry season | 4.61 [1.93 - 7.30] | 0.14 [0.07 - 0.20] |
| No. sample analysed for the presence of CSP | 1714 | 249 |
| Number of sporozoite positive | 59 | 8 |
| Overall SR % (95% CI) | 3.44% [2.52 - 4.68] | 3.21% [1.32 - 7.60] |
| SR wet season | 3.82% [2.56 - 5.66] | 3.59% [1.34 - 9.23] |
| SR dry season | 2.85% [1.71 - 4.72] | 1.85% [0.23 - 13.24] |
| Mean monthly EIR (weighted) | 4.57 [1.05 - 8.09] | 0.17 [0.01 - 0.33] |
| EIR wet season | 4.67[0.03 - 9.32] | 0.23 [0.01 - 0.45] |
| EIR dry season | 4.38 [0.69 - 8.07] | 0.07 [0.06 - 0.20] |
| Estimate annual EIR (weighted) | 54.85 [12.60 - 97.09] | 2.01 [0.12 - 3.91] |
| % EIR contribution (weighted) | 96.47% | 3.53% |

EIR: entomological inoculation rate; SR: sporozoite rate

Overall sporozoite rate varied across the study area with the highest rates observed in the southern zone (Table 2; OR: 1.88, [95% CI: 1.02-3.46]; p=0.044). In southern clusters, sporozoite rates for *An. funestus* s.l. was significantly higher than in northern clusters (OR: 2.33, [95% CI: 1.11-4.95]; p=0.028). The monthly sporozoite rate for *An. funestus* s.l. and *An. gambiae* s.l. fluctuated across the dry and wet seasons with slightly higher, but not significant, rates in the wet season (Table 3). *An. funestus* s.s. maintained malaria transmission across both seasons (sporozoite rates of 2.85% [95% CI: 1.71-4.72] and 3.82% [95% CI: 2.56-5.66], during the dry and wet seasons, respectively) while *An. gambiae* s.s. appeared to contribute to transmission mainly in the rainy season (sporozoite rates of 1.85% [95% CI: 0.23-13.24] and 3.59% [95% CI:1.34-9.23] during the dry and wet seasons, respectively) (Table 3).

In Misungwi district, malaria transmission occurs throughout the year. The average Entomological Inoculation Rate (EIR), weighted to account for the proportion of sampled *Anopheles* vectors processed for *Plasmodium* sporozoite infection, was 4.4 infective bites per house per month, approximately 53.3 per house per year, with variation in transmission intensities across the study area and seasons (Table 2 and 3). Overall, *An. funestus* s.s. was the major malaria vector responsible for 96.5% of total transmission (Table 3). Communities living in the southern part of the study area experienced significantly higher malaria transmission (EIR=9.6) compared to the northern zone (EIR=0.6) (Table 2). Monthly EIR was higher in the wet compared to the dry season, for both *An. funestus* s.l. (3.82 *vs*. 2.85) and *An. gambiae* s.l. (3.59 *vs*. 1.85; Table 3)

### *Anopheles* feeding and resting behaviours

A total of 1108 *Anopheles* vectors were sampled using four collection methods (CDC-LTs indoors, Furvela tent traps outdoors (Figure 3D), Prokopack aspirators indoors and outdoors), in 96 houses across 48 clusters between December 2018 and January 2019.

**Table 4.** Indoor and outdoor *Anopheles* feeding and resting behaviours and species composition.

| Outcome | CDC LT indoors | Furvela tent trap outdoors | Prokopack indoors | Prokopack outdoors |
| --- | --- | --- | --- | --- |
| Total HH/night collection | 96 | 96 | 96 | 96 |
| Total *Anopheles* vectors | 536 | 464 | 56 | 52 |
| Mean malaria vector | 5.6 [2.3 - 8.8] | 4.8 [2.9 - 6.7] | 0.6 [0.2 -0.9] | 0.5 [0.1 -1.0] |
| Proportion *An. gambiae* s.l. | 33.6% [14.4 -52.7] | 49.8% [32.2 - 67.4] | 51.8% [26.8 -76.7] | 71.2% [43.0 - 99.3] |
| Proportion *An. funestus* s.l. | 66.4% [47.3 -85.6] | 50.2% [32.6 -67.8] | 48.2% [23.3 - 73.2] | 28.8% [0.7 -57.0] |
| Proportion of *An. arabiensis* | 54.7% [25.5 - 81.0] | 51.8% [26.7 - 76.0] | 54.5% [12.5 - 91.0] | 89.5% [54.3 - 98.4] |
| Proportion of *An. gambiae* s.s. | 45.3% [19.0 -74.5] | 48.2% [24.0 -73.3] | 45.5% [9.0- 87.5] | 10.5% [1.6 - 45.7] |
| Proportion of *An. funestus* s.s. | 96.4% [91.7-98.5] | 99.4% [95.4-99.9] | 100% | 100% |
| Total *Anopheles* tested for CSP | 412 | 408 | 51 | 52 |
| Number of sporozoite positive | 7 | 7 | 2 | 0 |
| % SR | 1.7% [0.8 -3.5] | 1.7% [0.8 - 3.6] | 3.9% [1.0 - 14.4] | 0.0% |

CDC: Centers for Disease Control and Prevention; CSP: circumsporozoite protein; HH: household; LT: light trap; SR: sporozoite rate

As summarized in Table 4, the greatest proportions of *Anopheles* were sampled by indoor CDC-LTs (48.4%) and outdoor tent traps (41.9%). *An. arabiensis* and *An. gambiae* s.s. showed similar tendencies of feeding both indoors (54.7% and 45.3% collected in CDC-LTs, respectively) and outdoors (51.8% and 48.2% collected in tent traps, respectively) but *An. arabiensis* had a much stronger exophilic habit than *An. gambiae* s.s. (89.5% [95% CI: 54.3-98.4] *vs*. 10.5% [95% CI: 1.6-45.7] in Prokopack collections outdoors, respectively) (Table 4). *An. funestus* s.s. demonstrated similar behaviour to *An. gambiae* s.s., predominantly feeding indoors (CDC-LT collections) and outdoors (tent trap collections) (66.4% and 50.2%, respectively) and resting indoors (Prokopack collections) (48.2% [95% CI: 23.3-73.2]). Sporozoite rates were higher in samples collected indoors (range between 1.7% [95% CI: 0.8-3.5] and 3.9% [95% CI: 1.0-14.4]), compared to outdoors (range between 0% and 1.7% [95% CI: 0.8-3.6]) (Table 4). Malaria transmission both indoors and outdoors was solely due to *An. gambiae* s.s and *An. funestus* s.s.; none of the vectors collected resting outdoors were sporozoite positive.

### Phenotypic resistance and underlying molecular and metabolic resistance mechanisms

Wild populations of *An. funestus* s.l. and *An. gambiae* s.l. from across the study area were confirmed resistant to the diagnostic concentration of alpha-cypermethrin, with mean 30-minute knock-down ranging from 43.7% to 59.4% (Table 5). Similarly, both species were resistant to permethrin, with average 24-hour mortality ranging between 38.3% to 56.5% (Table 6). In general, levels of resistance to both pyrethroids were comparable between *An. gambiae* s.l. and *An. funestus* s.l., as well as between northern and southern study zones (Figure 2C and Tables 5 and 6).

Overall, the majority 92.2% (565/615) of *An. funestus* s.l mosquitoes tested in bioassays were confirmed by PCR as *An. funestus* s.s, with a small proportion of *An. parensis* (7.8%; 48/615). *An. gambiae* s.l. from bioassays that were tested for sibling species identification, were classified as similar proportions of *An. gambiae* s.s. (45.3%; 48/106) and *An. arabiensis* (54.7%; 58/106).

**Table 5.**
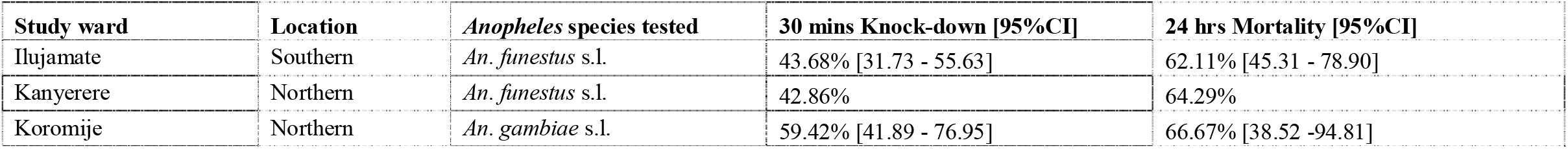
Average 30-minute knock-down and 24-hour mortality to the diagnostic dose of alpha-cypermethrin (CDC bottle bioassays: 12.5 µg/ml), among wild *Anopheles* species, collected from three study clusters in Misungwi, 2018.

**Table 6.** Average 60-minute knock-down and 24-hour mortality to the diagnostic dose of permethrin (WHO tube bioassays: 0.75%), among wild *Anopheles* species, collected from five study wards in Misungwi, 2018.

| Study ward | Location | *Anopheles* species tested | 60 mins Knock-down [95%CI] | 24 hrs Mortality [95%CI] |
| --- | --- | --- | --- | --- |
| Bulemeji | Northern | *An. gambiae* s.l. | 16.67% [9.09 - 28.57] | 38.33% [24.30 -52.37] |
| Idetemya | Northern | *An. gambiae* s.l. | 65.22% [47.32 -83.11] | 56.52%[35.21 - 75.67] |
| Ilujamate | Southern | *An. funestus* s.l. | 38.00% [26.65 -49.35] | 40% [28.39 -51.61] |
| Kanyerere | Northern | *An. gambiae* s.l. | 27.12% [13.92 - 40.32] | 38.98% [12.00 -65.97] |
| Mamaye | Northern | *An. gambiae* s.l. | 33.33% [19.30 - 47.37] | 45.00% [35.81 -54.19] |

Three hundred and twenty-two field collected *An. gambiae* s.l. were screened for the presence of L1014F-*kdr* and L1014S-*kdr* mutations. At the population level, homozygous L1014S-*kdr* was present in almost all *An. gambiae* s.s. individuals (98%; 90/92), with evidence for significant deviations from Hardy-Weinberg equilibrium (χ^2^=40.21; p<0.0001). No L1014S-*kdr* or L1014F-*kdr* were detected in any *An. arabiensis* tested (n=230); L1014F-*kdr* was not detected in any *An. gambiae* s.s. individuals.

Comparison of metabolic gene expression in *An. gambiae* s.s. collected from Mamaye ward (northern zone) demonstrated up-regulation of *CYP6M2* (fold change; FC=0.37 [95% CI: 0.20-0.43]), *CYP6P3* (FC=1.58 [95% CI: 0.89-2.07]), *CYP6P4* (FC=0.78 [95% CI: 0.46-1.11]) and *CYP9K1* (FC=1.58 [95% CI:1.19-4.80]).

Mean mortality 24 hours after exposure to the standard diagnostic dose of alpha-cypermethrin was predicted for *An. gambiae* s.l. in 2017 using a geospatial model. The model used data from WHO susceptibility tests conducted from 2005 to 2017 and incorporated associations between resistance and potential explanatory variables such as ITN coverage using three different machine learning approaches. Predicted mean mortality in *An. gambiae* s.l. for each 5 x 5 km square (Figure 2C) was high across Tanzania in 2017. Within Misungwi, the lowest mortalities / highest resistance occurred in the west and northwest.

## Discussion

Despite substantial gains achieved in malaria control across Tanzania over the past 20 years, attributable to improved quality and access to diagnostics and treatment and the widespread scale-up of LLINs and targeted IRS, localised transmission persists, especially in the Lake Region. Study findings demonstrate that *An. funestus* s.l. is becoming a dominant, efficient malaria vector species in Misungwi district, north-west Tanzania in an area with high coverage of standard pyrethroid LLINs and historical IRS activities. A similar phenomenon has recently been reported from south-eastern Tanzania [25]; however, our study indicates this shift in species composition may not be restricted to the south of the country.

Around Lake Victoria, species abundance and transmission intensity vary quite considerably spatially and temporally [28], with implications for the deployment of effective malaria vector control interventions. These heterogeneities likely reflect differences in climatic conditions such as rainfall and ecological settings, which support the breeding of particular vector species [29]. In our study, overall vector densities were significantly higher in villages located in the southern part of the study district compared to the northern clusters. *An. gambiae* s.s. and *An. arabiensis* occurred across Misungwi district, however, *An. gambiae* s.s. abundance was lower in the south and concentrated mostly in the north. By comparison, *An. funestus* s.s. was equally distributed throughout the district in sympatry with *An. arabiensis* and *An. gambiae* s.s., but found at the highest densities along shorelines and waterways feeding into Lake Victoria. The spatial variation of *Anopheles* sibling species may be explained by several factors linked to ecological features, including turbidity, water quality, relative humidity, temperature, vegetation type and/or socioeconomic parameters, such as ownership and usage of insecticide-based vector control measures and livestock density, as observed in previous studies conducted on the Kenyan side of Lake Victoria [33]. *An. gambiae* s.s. are known to breed in rain-dependent temporary habitats [34], while *An. funestus* s.s. and *An. arabiensis* can colonize large permanent aquatic habitats, some with large vegetation, in arid and highland areas [35,36]. Most residents in Misungwi district, especially in the southern clusters, traditionally stored rainwater for domestic purposes and animal husbandry in large, permanent man-made pools, locally called “Rambo”, which were filled throughout the year and could serve as potential breeding sites for *An. funestus* s.s. and *An. arabiensis*, even in the dry season; the higher density of *An. gambiae* s.s. during the rainy season is likely due to increased availability of temporary breeding sites [37,38]. In addition, agricultural practices such as irrigated rice paddies create diverse aquatic mosquito breeding habitats that could influence the co-existence and abundance of different vector species in the study area [39,40]. *An. funestus* s.l was collected in both seasons but peaked during the dry season, consistent with its ability to develop in habitats that can sustain desiccation [41]. Of concern, both *An. gambiae* s.l. and *An. funestus* s.l. malaria vector species were present during different seasons, favoured by distinct climatic and ecological conditions, sustaining malaria transmission throughout the region and the year. Of all mosquitoes sampled in this study, *Culex* species were the most abundant, contributing to 67.6% of the indoor host-seeking population. Previous studies in Tanzania have highlighted that *Culex* species, commonly referred to as “the house mosquito”, predominant in malaria-endemic communities [42], and when resistant to public health insecticides, can jeopardise community adherence to vector control interventions, due to perceived failure of these strategies [43,44].

This study estimated that each household could receive an average of more than 53 infective bites per year from both major vector species (*An. funestus* s.s. and *An. gambiae* s.s.) despite high coverage of LLINs. Comparably high EIRs have also been reported from other rural and peri-urban regions of East Africa, including south-central Tanzania [45], coastal Kenya [46] and southwest Ethiopia [47]. Furthermore, the annual EIR was ten times higher in villages located in the southern part of the study district compared to the north. In northern clusters, where *An. gambie* s.s. and *An. funestus* s.s co-existed, even though *An. gambiae* s.s. was present in very low numbers, these two species generally had equivalent *Plasmodium* infection rates. Both *An. gambiae* s.s. and *An. funestus* s.s. can be highly anthropophilic and endophilic, but the former species may be more aggressive and efficient vector in terms of malaria transmission, possibly due to host competition. In the southern study clusters, malaria transmission was almost exclusively mediated by *An. funestus* s.s. Only a single *P. falciparum*-infected *An. arabiensis* was collected during the study which might be explained by its highly opportunistic behaviour, feeding on both animals and humans; in the absence of the latter host it can display strongly zoophilic feeding preferences for livestock, of which close to half of the households owned [48,49]. This species is also known for its more exophilic tendencies compared to *An. gambiae* s.s. [49,50] and can easily adapt and feed outdoors in response to insecticidal interventions [51], especially when human or animal populations are available outside [12]. Our indoor and outdoor collections generally support these behavioural assumptions, with both *An. gambiae* s.s. and *An. funestus* s.s. sampled in similar proportions across different traps, with the exception of *An. gambiae* s.s., which was found at very low densities resting outdoors. The occurrence of highly endophilic and anthropophagic vectors such as *An. funestus* s.s. host-seeking or resting outdoors could be linked to behavioural divergence among vector populations and/or chromosomal inversion polymorphisms [52,53], as well as human behavioural changes [54]. However, in our study, more sporozoite-harbouring mosquitoes were collected in houses, suggesting ongoing malaria transmission is still occurring inside, despite high intervention coverage. The strongly exophilic behaviour of *An. arabiensis* indicated that LLINs and IRS in Misungwi district may have minimal effect against this species; although its relative importance in local malaria transmission appears diminished. Moreover, the presence of infected *An. gambiae* s.s. and *An. funestus* s.s. outdoors, coupled with a degree of exophagic behaviour, suggests that additional control tools targeting outdoor vector populations may warrant consideration in the study area [55].

All three major vector species (*An. gambiae* s.s., *An. arabiensis* and *An. funestus* s.s.), displayed low levels of susceptibility to alpha-cypermethrin and permethrin, primarily due to selection pressure from prolonged use of pyrethroid-based LLINs and likely enhanced by concurrent agricultural pesticide application [56-58]. Study findings align with others in the Lake Zone and across Tanzania, demonstrating low mortality among vector populations to the diagnostic doses of pyrethroids [9,57,59]. Data collected across Africa indicated that previously pyrethroid resistance was higher in east African populations of *An. funestus* s.l. compared to east African populations of *An. gambiae* s.l. up to 2017, but this difference was not detected in Misungwi in 2018-19 [60,61]. It is noteworthy that our bioassays presented higher levels of resistance in *An. gambiae* s.l. in comparison to those shown in our map produced by geospatial models of phenotypic resistance to alpha-cypermethrin for the year 2017, suggesting pyrethroid resistance may be increasing in *An. gambiae* s.l. populations in the region. In Misungwi district, population-level frequency of the L1014S-*kdr* mutation was practically fixed in *An. gambiae* s.s., while *CYP6M2, CYP6P3, CYP6P4* and *CYP9K1* were modestly upregulated by comparison to reports from West Africa [62-64]. These results indicate that both target site and metabolic mechanisms may be driving pyrethroid resistance in *An. gambiae* s.s. in this study area. However, further investigation is necessary to identify resistance mechanisms specific to these field populations for prospective monitoring and to improve our understanding of the specificity of resistance mechanisms to individual interventions and the likelihood of selecting for cross-resistance between active ingredients [62-64]. Importantly, intense insecticide resistance may partially explain the persistent malaria transmission in Misungwi district, highlighting the urgent need for novel vector control tools, containing different insecticide classes and combinations. This study was undertaken prior to the phase III evaluation of novel bi-treated LLINs containing a pyrethroid and either a pyrrole (chlorfenapyr), a synergist (piperonyl butoxide; PBO) or a juvenile growth hormone inhibitor (pyriproxyfen; PPF) [31], which may have the potential to control malaria transmitted by pyrethroid-resistant vector species.

This study was conducted to characterize baseline vector population bionomics and malaria transmission dynamics in Misungwi district, with some limitations. Because mosquito collections spanned five months of the year, encompassing the short rainy season (October to December 2018) and part of the dry season (August and September 2018), vector densities, sibling species composition and sporozoite rates reported in this study may not be representative of the annual variation in this area. Additional studies are ongoing investigating in-depth the association between vector spatial distributions and key ecological indices, and to identify insecticide resistance mechanisms in *An. funestus* s.l., which at the time of study design, was not anticipated to emerge as the major vector species in Misungwi district.

## Conclusion

In Misungwi district, North-West Tanzania, *An. funestus* s.s. is the leading malaria vector species, predominating in southern villages of the study site, across dry and wet seasons. *An. gambiae* s.s was present in much lower densities, concentrated mostly in the north during the wet season, potential driving malaria epidemics. Annual EIR was high, despite high LLIN usage, but variable within a small geographical area, influenced by vector species diversity and bionomics, with serious epidemiological implications for malaria control. *An. gambiae* s.s., *An. arabiensis* and *An. funestus* s.s. were found similarly resistant to pyrethroids, with high frequencies of target site alleles and overexpression of detoxification genes identified in *An. gambiae* s.s. Study findings highlight the urgent need for novel vector controls strategies, which incorporate new chemical classes, to control malaria transmitted by pyrethroid-resistant vector populations and sustain gains in malaria control across the Lake Region.

## Methods

### Study area characteristics

The study was carried in Misungwi district (latitude 2.85000 S, longitude 33.08333 E) in North-West Tanzania on the southern shore of Lake Victoria (Figure 1A). Misungwi lies at an altitude of 1,150 meters above sea level, with a population of approximately 351,607 according to the National population and housing census of 2012 [65]. The district experiences a dry season typically between June and September and two annual rainy seasons; the long-rainy season between February and May and a short-rainy season between November and December with average annual rainfall ranging between 0.5 and 58.8 mm. The district is geographically divided into two main agro-ecological zones (northern and southern zones), based on the vegetation land cover and amount of rainfall. The local communities practice rice, millet and cotton farming, domestic animal rearing, fishing and have small-scale businesses as sources of income and food. In preparation for the CRT, the study area was sub-divided into 86 clusters, containing 72 villages made up of 453 hamlets from 17 wards (Figure 2). Detailed criteria and methodology used for cluster formation is described elsewhere [31].

Across Misungwi district, a typical compound was comprised of a main house and cattle shed. Houses were generally constructed from both modern and traditional materials and most houses had eave spaces (an opening between the wall and the roof for ventilation) (Figure 3A and B). The area experiences moderate to high malaria transmission and malaria incidence peaks shortly after the rainy seasons [66]. Recent studies conducted in 2010 and 2017 reported a malaria prevalence of 51.3% across all age groups and 46.3% in school children (7-14 years). LLINs mainly Olyset® and PermaNet® 2.0 obtained through national bed net distribution campaigns have been the primary malaria control method in the study area [4,67] and IRS was last conducted in this region in 2015. A preliminary survey carried out by our study team in 2018 found *An. gambiae* s.s. *An. arabiensis* and *An. funestus* s.s. as the predominant malaria vector species in the study area.

### Environmental characteristics of northern and southern clusters

To characterize the study area with regards to climatic and environmental conditions, high spatial resolution satellite remote sensing and other geospatial data were downloaded in raster (i.e. gridded) format from publicly available data sources and processed using ArcGIS 10.5.1 (ESRI, Redlands, USA). Data on eight bioclimatic variables at 1 km^2^ spatial resolution, representing averages for the years 1970-2000, were obtained from the WorldClim 2 database [68]: annual mean temperature (Bio1), temperature seasonality (Bio4), maximum and minimum temperature of the warmest month (Bio 5 and 6), annual precipitation (Bio12), precipitation of the wettest and driest months (Bio13 and 14), and precipitation seasonality (Bio15). Global elevation data were obtained for the study area from NASA’s Shuttle Radar Topography Mission (SRTM) 4.1 at 90-meter spatial resolution [69]. Global landcover data were obtained from the European Space Agency GlobCover 2009 project, available at 300-meter spatial resolution (© ESA 2010 and UCLouvain; http://due.esrin.esa.int/page_globcover.php). These data identify 22 landcover types, of which 12 were identified in the study area. Zonal mean statistics for the northern and southern clusters were calculated for each bioclimatic variable and elevation using the spatial analyst toolbox in ArcGIS; cluster means were then averaged for each zone (Supplementary Table 1). The proportion of cells within each of the northern and southern zones that were classified as different landcover types were similarly calculated; Supplementary Table 1 shows the results for the five dominant landcover types that were present in the study area [mosaic vegetation (i.e. grassland/shrubland/forest: 50-70% / cropland: 20-50%), herbaceous vegetation (i.e. grassland/savannas), shrubland, broadleaf deciduous forest/woodland, and grassland or woody vegetation on regularly flooded or waterlogged soil], which represent 98.9% and 97.5% of the total area of the northern and southern clusters, respectively. While the available data sources, and hence these estimates, are derived from time periods prior to the study period, we assume that the estimates accurately reflect relative differences in climatic and environmental conditions across the study area.

### Indoor and outdoor entomological surveillance

Two cross-sectional entomological field surveys were conducted between August and December 2018 in all 86 clusters, using CDC-LTs (John W Hock Company, USA). Eight households were randomly selected from a census list of households generated during baseline enumeration. CDC-LTs were hung next to the feet of an occupant sleeping under an ITN/untreated net (about 1m from the ground), between 19:00 and 7:00. A questionnaire was administered to the head of the household to gather information related to the house structure (type of wall and roofing materials, windows screening, number of rooms, number of sleeping places, presence of eaves), and coverage and usage of LLINs/untreated nets. Direct observation was also used during data collection to validate participant answers.

Assessment of *Anopheles* vector feeding and resting behaviours indoors and outdoors was undertaken in 48 clusters between December 2018 and January 2019. Two households per cluster were randomly selected, and each house was installed with a CDC-LT indoors and an occupied Furvela tent trap outdoors [70]. Both Furvela and CDC-LTs were switched on at 19:00 and off at 7:00. Indoor and outdoor resting adult *Anopheles* were collected from the same houses using a 12 voltage battery-powered Prokopack aspirator [71], and manual mouth aspirators [72,73]. Systematic sampling of adult resting mosquitoes on the walls, roofs and floors were conducted for up to three minutes depending on the size of the room. Outdoor collections were performed from potential resting sites around the house, such as open resting structures, cow sheds and pit latrines.

### Insecticide resistance testing

Insecticide resistance profiles of wild populations of *An. gambiae* s.l and *An. funestus* s.l were assessed in six clusters selected on the basis of high *Anopheles* population densities. Adult female *Anopheles* were collected resting indoors using both Prokopack and mouth aspirators [71]. Mosquitoes were separated by species complex and supplied with 10% glucose solution for 72 hours to allow digestion of blood meal, prior to bioassay testing with permethrin and alpha-cypermethrin. In WHO tube assays, 20-25 gravid female *An. gambiae* s.l. *or An. funestus* s.l. of unknown age were exposed to 0.75% permethrin for 60 minutes [74]. In CDC bottle bioassays, 20-25 gravid female *An. gambiae* s.l. or *An. funestus* s.l. of unknown age were exposed to 12.5 μg/ml alpha-cypermethrin for 30 minutes [75]. For both assays, knock-down was recorded at the end of the diagnostic exposure time (30 or 60 minutes after exposure, for CDC or WHO bioassays, respectively), and final mortality was scored after 24 hours. All mosquitoes tested in bioassays were stored individually for sibling species identification.

### Mosquito processing, species identification and sporozoite detection

Adult female mosquitoes collected from the cross-sectional surveys, resistance bioassays and behaviour study were sorted and identified based on their morphology, separating *An. gambiae* s.l. from *An. funestus* s.l. and from other genera according to Gillies and Coetzee [76]. At least three female *An. gambiae* s.l. and three *An. funestus* s.l. per household/ per collection method was analysed for presence of *Plasmodium falciparum* CSP using enzyme-linked immunosorbent assay (CSP-ELISA) [77]. All CSP positive samples and a sub-sample of CSP negative *An. gambiae* s.l. and *An. funestus* s.l. from the cross-sectional surveys, resistance bioassays and behaviour study, were randomly picked per house and tested for species identification. DNA was extracted from legs/wings and TaqMan assays were performed to distinguish sibling species in *An. gambiae* [78] or *An. funestus* complexes [79].

### Identification of insecticide resistance mechanisms

A subsample of *An. gambiae* s.s. and *An. arabiensis* were genotyped for L1014F-*kdr* and L1014S-*kdr* mutations associated with pyrethroid and DDT resistance, using TaqMan PCR assays. [80]. Blood-fed indoor resting female adult mosquitoes (F0s) were collected using mouth aspirators from three wards, sampled for phenotypic resistance testing. Mosquitoes were held for 3-4 days to allow for blood meal digestion. Individual *An. gambiae* s.l were placed into Eppendorf tubes containing moist filter papers and forced to lay eggs, as previously described [23]. The first 3-5 emerged F1 adults from each parent were stored individually in RNAlater® and preserved at-20°C for gene expression analysis. Expression profiles for metabolic detoxification genes in a sub-sample of 250-300 F1 wild-caught female *An. gambiae* s.s. mosquitoes were determined using quantitative reverse transcriptase PCR (qRT-PCR) [81,82]. Individual mosquitoes were first tested for species identification, and only mosquitoes identified as *An. gambiae s*.*s*. were analyzed. A minimum of 5 pools of 10 *An. gambiae* s.s were analysed for *CYP6M2, CYP6P3, CYP6P4* and *CYP9K1* gene expression [81].

### Data analysis

Field data were entered into an Open Data Kit (ODK) form. Data analysis was performed using Stata/IC 15.1 (Stata Corp., College Station, USA) [83]. Mean *Anopheles* caught per night per household, sporozoite rate and their 95% confidence intervals (CIs) were estimated according to study zones (North or South), season (wet or dry) and *Anopheles* species. *Anopheles* vector population density and entomological inoculation rate (EIR) were analysed and compared between study zones and seasons using multilevel negative binomial regression taking into account clustering effect. The EIR was calculated at household level as the average number of CSP-ELISA positive mosquitoes per night and was weighted to account for the proportion of collected *Anopheles* processed for CSP-ELISA. Logistic regression was used to compare sporozoite rates between the two study zones and seasons. The proportion of households with at least one LLIN was computed by dividing total nets observed and recorded by total households surveyed. Net access was estimated from the proportion of households with enough LLINs over total sleeping places. Unimproved houses were classified as houses with open eaves, unscreened windows, and were constructed from traditional low-quality materials such as a thatched roof, mud and non-plastered walls. Improved houses had closed eaves, with mosquito proof mesh on the windows, and were built with improved modern materials such as an iron sheet as a roof, brick/blocks, with plastered walls.

For resistance phenotyping, mean percentage knock-down/mortality post-exposure was calculated and interpreted following the WHO and CDC criteria [74,75]. Susceptibility thresholds were considered at the diagnostic time (24 hours and 30 minutes post-exposure for WHO and CDC bioassays, respectively). Mean mosquito mortality between 98 and 100% indicated full insecticide susceptibility, 90-97% showed suspected resistance that needed confirmation and less than 90% indicated confirmed resistance [74,75]. When control mortality was between 5-20%, results were corrected using Abbott’s formula [74]. If the control mortality was ≥ 20%, the test was discarded [74].

Gene expression and fold-change, relative to the susceptible laboratory strain *An. Gambiae* s.s. Kisumu were calculated according to the 2-ΔΔCq method [84] after standardisation with housekeeping genes (elongation factor; *EF* and 40S ribosomal protein S7; *RPS7*).

To estimate spatial and temporal trends in pyrethroid resistance using the available field data, which is sparse and has a heterogeneous distribution, a total of 6,423 observations of mortality from WHO susceptibility tests from 2005-2017 were used to inform a Bayesian geostatistical ensemble model. The model was also informed by a suite of 111 potential explanatory variables describing potential drivers of selection for resistance such as ITN coverage and produced estimates of mean mortality in *An. gambiae* s.l. for two regions of Africa [61]. Here we present the results for alpha-cypermethrin resistance in Misungwi in 2017.

### Ethical approval and consent to participate

This study is part of an ongoing RCT in Misungwi (clinical trial registration: <u>NCT03554616</u>) which obtained ethical clearance from the National Institute for Medical Research (NIMR), Tanzania (NIMR/HQ/R.8a/Vol. IX/2743) and the London School of Tropical Medicine and Hygiene, United Kingdom (LSHTM ethics ref: 14952) [31]. All study procedures were performed in accordance with relevant guidelines and regulations. Prior to study initiation, community consent was sought from village leaders and written, informed consent was obtained from the heads of all households selected for participation. Study information, including the study purpose, risks and benefits, was provided to participants in Swahili.

### Data availability

The data sets generated and/or analysed during the current study are not public but are available from the corresponding author on reasonable request

## Acknowledgments

We thank the entomology technicians for their support in the field work and Misungwi District residents for their acceptance and cooperation throughout the study. This study was funded by the Department for International Development, the UK Medical Research Council, the Wellcome Trust and the Department of Health and Social Care (#MR/R006040/1).

## Author contributions

NP, FWM, MR, MAK, JFM and AM wrote the main study protocol and design the study. NSM, NP, LAM, PH, CM performed data analysis. GI, BS and RB performed the molecular analysis. JM and NSM supervised the study data collections. NSM wrote the initial draft of the manuscript, which was revised by NP and LAM. All authors read and approved the final manuscript.

